# CNV-BAC: Copy Number Variation Detection in Bacterial Circular Genome

**DOI:** 10.1101/2019.12.24.887992

**Authors:** Linjie Wu, Han Wang, Yuchao Xia, Ruibin Xi

## Abstract

**Motivation:** Whole genome sequencing (WGS) is widely used for copy number variation (CNV) detection. However, for most bacteria, their circular genome structure and high replication rate make reads more enriched near the replication origin. CNV detection based on read depth could be seriously influenced by such replication bias.

**Results:** We show that the replication bias is widespread using ~200 bacterial WGS data. We develop CNV-BAC that can properly normalize the replication bias as well as other known biases in bacterial WGS data and can accurately detect CNVs. Simulation and real data analysis show that CNV-BAC achieves the best performance in CNV detection compared with available algorithms.

**Availability and implementation:** CNV-BAC is available at https://github.com/LinjieWu/CNV-BAC.

## 1 Introduction

Whole genome sequencing (WGS) has been widely used to study genomic variations in many different organisms including bacteria. Copy number variation (CNV), consisting of gains or losses of DNA segments, is an important class of genomic variations. CNVs are often detected using the read depth (RD) information of WGS. However, other than copy number states, RD can be easily influenced by factors such as GC-content and the mappability of the genomic regions. Hence, proper normalization needs to be performed before CNV detection (Hansen, et al., 2010). For almost all bacteria, their single circular chromosome replicates bidirectionally from a single fixed origin towards a single terminus (Wang and Levin, 2009). Therefore, during replication, a bacterium should have two copies of DNA sequences that has been replicated and only one copy of DNA sequences yet to be replicated. Since WGS of bacteria is performed using millions of bacterial cells and cells may be at various cell cycle stages, regions closer to the replication origin should have higher RD than regions closer to the replication terminus. This replication bias could seriously influence the CNV detection if not properly accounted for. In this paper, we use ~200 bacteria WGS data from 3 studies to show that this replication bias is widespread. We develop a new algorithm called CNV-BAC that can properly normalize the replication bias and can accurately detect CNVs in bacterial genomes. Comprehensive simulation and real data analysis show that CNV-BAC achieves best performance in detecting CNVs in bacterial genomes.

## 2 Methods

In the normalization step, CNV-BAC first applies the normalization procedure of BIC-seq2 (Xi, et al., 2016) to normalize against GC-content and the mappability. For each bin *i*, let *O*_*i*_ be the observed RD, *E*_*i*_ be the expected RD given by BIC-seq2 and *d*_*i*_ be the distance of the bin *i* to the replication origin. CNV-BAC uses Gaussian or Poisson Model to model the dependence of the observed RD *O*_*i*_ on the distance to the replication origin *d*_*i*_ and the expected RD *E*_*i*_ (see Supplementary Text). After normalization, CNV-BAC gives a new expected RD *Ê*_*i*_ and uses the segmentation algorithm of BIC-seq2 to call CNVs. See Supplementary text for more details of the model and implementation.

## 3 Result

### 3.1 Real Data Analysis

We consider three bacterial sequencing datasets, (1) 55 *Escherichia coli* (*E. coli*) MDS42 cells (Horinouchi, et al., 2017), (2) 85 *Staphylococcus aureus* (*S. aureus*) strains (Copin, et al., 2019), and (3) 50 *Lactobacillus casei Zhang* (*L. casei Zhang*) (Wang, et al., 2017). After mapping, we calculate the Spearman’s rank correlation between the RD *O*_*i*_ and the bin’s distance to the replication origin *d*_*i*_. For all three datasets, as predicted from the mechanism of the replication bias, most correlations are less than zero (Figure 1A). We further calculate the Spearman’s correlation between *d*_*i*_ and the log2 copy ratio *Y*_*i*_ = log_2_(*O*_*i*_/*E*_*i*_) given by BIC-seq2. Interestingly, possibly because BIC-seq2 has removed other sources of biases, the negative correlation between *d*_*i*_ and *Y*_*i*_ becomes more prominent (Figure 1A). A number of bacteria even have correlation close to −1 (e.g. −0.945 for *S. aureus* genome SRR8074817). In comparison, after CNV-BAC’s normalization, the correlations between *d*_*i*_ and the log2 copy ratio *Ŷ*_*i*_= log_2_(*O*_*i*_/*Ê*_*i*_) are very close to zero (Figure 1A).

**Fig. 1.**
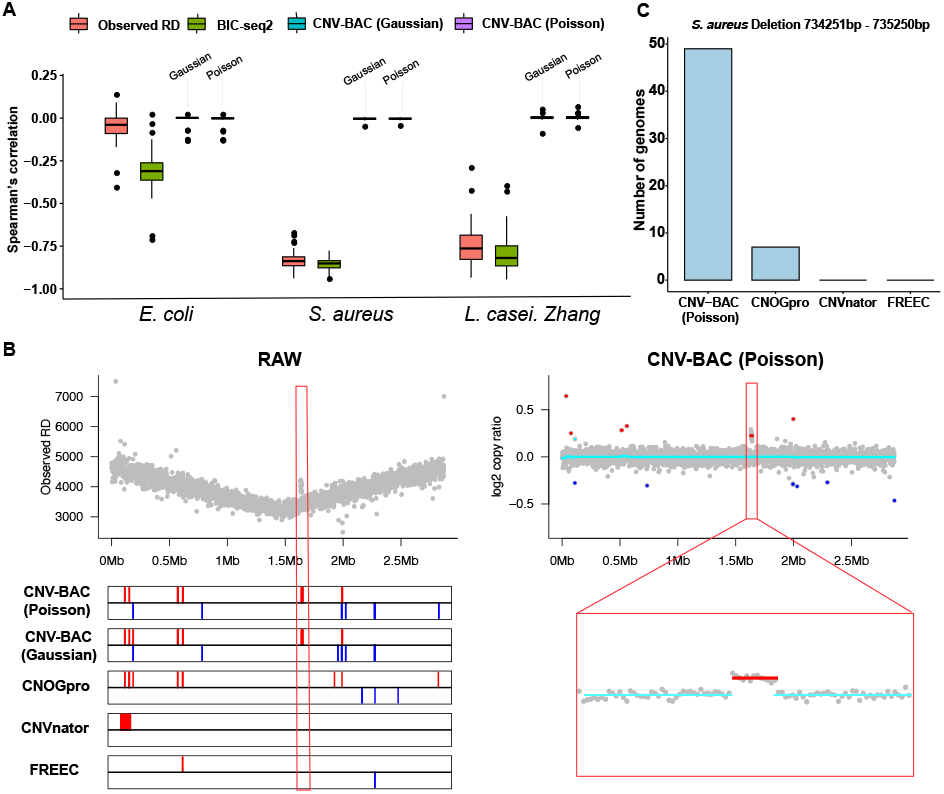
CNV detection for real data. (A) The boxplot of Spearman’s rank correlations of the observed RD, the log2 copy ratios given by BIC-seq2 and CNV-BAC with the distance to replication origin. (B) The RD profile and detected CNVs of *S. aureus* SRR8074817. Top left: the raw RD; Bottom left: CNVs detected by different algorithms (bars in red are gains and bars in blue are deletions); Top right: the log2 copy ratio given by CNV-BAC based on the Poisson model (red/blue lines show the significant log2 copy ratio for duplication/deletion segments and cyan lines are normal segments); Bottom right: The zoom-in view of the apparent duplication missed by other algorithms. (C) The number of *S. aureus* genomes that are detected to have a deletion (734251bp - 735250bp) by different methods.

Figure 1B shows the *S. aureus* genome SRR8074817. Its raw data demonstrates very strong replication bias (top left panel) and this bias disappears after CNV-BAC normalization (right panel). In addition, this genome has a very clear copy number gain (marked in red box). Since the gain’s signal is masked by the replication bias, available algorithms such as CNOGpro (Brynildsrud, et al., 2015), CNVnator (Abyzov, et al., 2011) and Control-FREEC (Boeva, et al., 2012) fail to detect the gain. In comparison, CNV-BAC successfully detects the gain. Supplementary Figure S1 shows more of similar examples. The distribution of CNV numbers detected by these algorithms are shown in Supplementary Figure S2. Overall, CNVnator is the most conservative and it only detects 0.7 CNVs per sample on average. FREEC is also conservative, detecting 2.6 CNVs per sample. We further use the normalization data of CNV-BAC to estimate the false discovery rate (FDR) of available algorithms (Supplementary Text), CNVnator has the lowest FDR, followed by FREEC and CNOGpro (average FDR 0.27, 0.40 and 0.85 respectively). Interestingly, in *S. aureus*, CNV-BAC identifies a deletion (73425bp - 735250bp) occurred only in the high-risk strains of *S. aureus* (Supplementary Figure S3). In total, CNV-BAC identifies 49 strains having this deletion, and the other algorithms only identify less than 7 strains having this deletion (Figure 1C). Similarly, CNV-BAC finds 19 strains of *L. case. Zhang* having a common deletion, and other algorithms only find 1 strain (Supplementary Figure S4).

### 3.2 Simulation Analysis

We use simulation to compare CNV-BAC with available algorithms including BIC-seq2, CNVnator, Control-FREEC and CNOGpro. See Supplementary Text for detailed simulation setup (Supplementary Figure S5-S6). Overall, CNV-BAC achieves the best sensitivity and specificity in most simulation scenarios. Largely speaking, smaller events are harder to detect for all algorithms. As the sequencing depth increases, the powers tend to be larger for all algorithms. The power of CNV-BAC keeps largely the same for different levels of replication bias for most scenarios. In terms of FDR, CNV-BAC and FREEC have the lowest false positives and largely keeps constant when the replication bias increases. The false positives of other algorithms tend to be high when the replication bias is high. As expected, the false positive duplications are mostly located near the replication origin, and the false positive deletions are mostly located near the replication terminus (Supplementary Figure S7-S15).

## References

Abyzov, A., et al. CNVnator: An approach to discover, genotype, and characterize typical and atypical CNVs from family and population genome sequencing. Genome Research 2011;21(6):974–984.

Boeva, V., et al. Control-FREEC: a tool for assessing copy number and allelic content using next-generation sequencing data. Bioinformatics 2012;28(3):423–425.

Brynildsrud, O., Snipen, L.G. and Bohlin, J. CNOGpro: detection and quantification of CNVs in prokaryotic whole-genome sequencing data. Bioinformatics 2015;31(11):1708–1715.

Copin, R., et al. Sequential evolution of virulence and resistance during clonal spread of community-acquired methicillin-resistant Staphylococcus aureus. Proceedings of the National Academy of Sciences of the United States of America 2019;116(5):1745–1754.

Hansen, K.D., Brenner, S.E. and Dudoit, S. Biases in Illumina transcriptome sequencing caused by random hexamer priming. Nucleic Acids Res 2010;38(12).

Horinouchi, T., et al. Prediction of Cross-resistance and Collateral Sensitivity by Gene Expression profiles and Genomic Mutations. Scientific Reports 2017;7.

Wang, J.C., et al. Genome adaptive evolution of Lactobacillus casei under long-term antibiotic selection pressures. BMC Genomics 2017;18.

Wang, J.D. and Levin, P.A. OPINION Metabolism, cell growth and the bacterial cell cycle. Nat Rev Microbiol 2009;7(11):822–827.

Xi, R.B., et al. Copy number analysis of whole-genome data using BIC-seq2 and its application to detection of cancer susceptibility variants. Nucleic Acids Res 2016;44(13):6274–6286.

