## Supplementary text for "CNV-BAC: Copy Number Variation Detection in Bacterial Circular Genome"

#### 1. Model and Algorithm

##### 1.1 Normalization Model

**Model**

In the normalization step, CNV-BAC first applies the normalization procedure of BIC-seq2 (Xi, et al., 2016) to normalize against GC-content and the mappability. For each variable-sized bin $i$ as defined in the BIC-seq2 paper (by default, we use 1000bp variable-sized bin), BIC-seq2 gives an observed number of reads $O_i$ and an expected number of reads $E_i$. Suppose that $d_i$ is the distance of the bin $i$ to the replication origin. The replication origin of the bacteria genomes can be found in the Doric database (Luo and Gao, 2019). We use two different approaches to model the dependence of the observed read count $O_i$ on the distance to the replication origin $d_i$. In the first approach, we model the random variable $Y_i = \log_2(O_i/E_i)$ as

$$Y_i = f(d_i) + \epsilon_i,$$

where $f$ is an unknown smooth function and $\epsilon_i$ is a Gaussian noise. In the second approach, we model $O_i$ as a Poisson random variable with mean $\lambda_i$ and assume that

$$\log(\lambda_i) = \beta_0 + \beta_1 E_i + g(d_i).$$

where the parameters $\beta_0$ and $\beta_1$ and the smooth function $g$ are to be estimated. In both approaches, the unknown functions $f$ and $g$ are represented as linear combination of spline base functions. We use the R package *mgcv* to estimate the unknown parameters and functions. In the first approach, assume that $\hat{f}$ is the estimate of $f$. The new expected read count in bin $i$ is defined as $\hat{E}_i = 2^{\log_2 E_i + \hat{f}(d_i)}$. In the second approach, assume that $\hat{\beta}_0$, $\hat{\beta}_1$ and $\hat{g}$ are the estimates of the unknown parameters and function. The new expected read count is defined as $\hat{E}_i = exp\ (\hat{\beta}_0 + \hat{\beta}_1 E_i + \hat{g}(d_i))$.

**Estimation Procedure**

CNV-BAC uses the following three steps to estimate the unknown functions and parameters.

a. Estimate parameters $f(\cdot)$ or $\beta_0$ , $\beta_1$ and functions $g(\cdot)$ by the gam function using all bins.
b. Calculate residuals of fitted model in step a. Let $Q_{0.75}^{res}$ and $Q_{0.25}^{res}$ be the 0.75th quantile and 0.25th quantile of the residuals and $IQR^{res} = Q_{0.75}^{res} - Q_{0.25}^{res}$. Filter the bins whose residuals are larger than $Q_{0.75}^{res} + 1.5 * IQR^{res}$ or less than $Q_{0.25}^{res} - 1.5 * IQR^{res}$. Use the remaining bins to estimate $\hat{f}$ or $\hat{\beta}_0$, $\hat{\beta}_1$ and $\hat{g}$.
c. Calculate predicted log2 copy ratios using estimations from step b. Let $Q_{0.75}^{CR}$ and $Q_{0.25}^{CR}$ be the 0.75th quantile and 0.25th quantile of the predicted log2 copy ratios and $IQR^{CR} = Q_{0.75}^{CR} - Q_{0.25}^{CR}$. Filter the bins whose log2 copy ratios are larger than $Q_{0.75}^{CR} + 1.5 * IQR^{CR}$ or less than $Q_{0.25}^{CR} - 1.5 * IQR^{CR}$. Use the remaining bins to get the final estimate $\hat{f}$ or $\hat{\beta}_0$, $\hat{\beta}_1$ and $\hat{g}$.

**CNV detection**

We provide the observed read count $O_i$ and the expected read count $\hat{E}_i$ to the segmentation algorithm of BIC-seq2 to segment the data. Then, for each segment $s$ given by BIC-seq2, let $O_s$ and $\hat{E}_s$ be the sum of the observed read counts and the expected read count of bins in the segment $s$, and $K_s$ be the number of bins in the segment $s$. Define $r_s = \log_2(O_s/\hat{E}_s)$ be the log2 copy ratio of this segment. Then, we bootstrap from the bins $B$ times to give the p-value to this segment. In the $b$th bootstrap, we randomly select $K_s$ bins and calculate the total observed read count $O_b$ and $\hat{E}_b$ of these $K_s$ bins. Let $r_b = \log_2(O_b/\hat{E}_b)$. Assume that $\mu$ and $\sigma$ be the sample mean and sample standard deviation of these $B$ log2 copy ratios $r_b$. CNV-BAC gives the p-value as $1 - 2\Phi(|r_s - \mu|/\sigma)$, where $\Phi$ is the cumulative distribution function of the standard normal distribution. Segments with Benjamini-Hochberg adjusted p-values smaller than 0.01 and absolute values of log2 copy ratio larger than 0.2 were selected as putative CNVs. For Control-FREEC, CNVnator and CNOGpro, we used parameters recommended by their manuals.

#### 2. Data analysis

##### 2.1 Simulation

To generate simulated sequencing data, we first generate a template genome by placing 9 duplications and 9 deletions of size 5 kb, 10 kb and 500 kb at different locations to the reference genome NC_020518 of E. coli (Supplementary Table S1). Then, we generate two 500X sequencing data from the template genome and from NC_020518. We consider 48 different scenarios (explained below) and randomly sample 100 data from these two 500X data for each scenario (Supplementary Figure S6).

We consider four levels of dependence on the distance to the replication origin measured by the simulating parameter $\beta$. The larger the parameter $\beta$ is, the more the dependence is (Supplementary Figure S7). We use uneven subsampling to simulate the influence of replication on the RD. Since in real data, CNVs may not exist in all sequenced bacteria, we also mix the reads generated from the template genome and from NC_020518. The proportions of reads from the template genome are set as 100%, 70%, 50% and 30% (we call these proportions the purities). Lastly, we consider three different coverage levels (10X, 50X and 100X). For each of the 48 simulating scenarios (4 levels of replication dependence × 4 levels of purities × 3 levels of coverage), we randomly subsample 100 data for each scenario, and we get 4800 sequencing data in total. The simulated reads were mapped to NC_020518 by BWA (Li and Durbin, 2009).

We use the overlap of detected CNVs and designed CNVs to infer true and false discovered CNVs for each sample. For a putative CNV detected by a given method, if the overlapping region is more than half of a designed CNV, this designed CNV is claimed to be detected in the current sample. If the overlapping region with any of the designed CNVs is less than half of the discovered CNV, we define it to be a false positive. Supplementary Figure S7-S14 show the sensitivity and FDR of the CNV detection algorithms. The false positives of other algorithms tend to be high when the replication bias is high. As expected, the false positive duplications of other algorithms are mostly located near the replication origin, and the false positive deletions are mostly located near the replication terminus (Supplementary Figure S15).

##### 2.2 Real Data Analysis

The original *E. coli* data contains 58 genomes, the *S. aureus* data contains 92 genomes, and *L. casei Zhang* data contained 50 genomes. We use BWA to map these sequencing data to reference genomes NC_020518.1, NC_007793.1 and CP001084.2 for *E. coli*, *S. aureus* and *L. casei Zhang*, respectively. We use the normalization result of BIC-seq2 and calculated log2 copy ratio for each bin. The noise level was defined as the mean of square difference of log2 copy number ratio between adjacent bins:

$$noise\ level = \frac{\sum_{i=1}^{n-1}(r_{i+1} - r_i)^2}{n-1},$$

where $r_i$ is the log2 copy number ratio of the $i$-th bin. For three bacterial datasets, we remove samples with high noise level (larger than 0.02). After quality control, the *E. coli* data contains 55 E. coli MDS42 cells cultured under 11 environmental stresses. The *S. aureus* has 85 *S. aureus* strains that are methicillin resistant. The *L. casei Zhang* data contains 50 *L. casei Zhang* strains cultured in three different culture media over 2000 generations. Average coverages for *E. coli*, *S. aureus* and *L. casei Zhang* datasets are 300X, 450X and 200X.

We use the following procedure to estimate the FDRs of other available algorithms. Given a candidate CNV detected by an algorithm. We first find all bins that overlap with this candidate CNV and calculate the total observed read count $O$ and total expected read counts $\hat{E}$ given by CNV-BAC in these bins. Then, we use the bootstrap method as described in the CNV Detection section to determine a p-value of the candidate CNV. If the p-value is greater than 0.05, we classify this CNV as a false positive.

### Supplementary Table and Figures

Table S1: Designed copy number variation regions for simulation data.

| Duplications | | | | Deletions | | | |
|---|---|---|---|---|---|---|---|
| Start | End | Length | Distance to replication origin | Start | End | Length | Distance to replication origin |
| 600001 | 605000 | 5000 | 1221472 | 20001 | 25000 | 5000 | 641472 |
| 1100001 | 1105000 | 5000 | 1721472 | 1700001 | 17050000 | 5000 | 1654347 |
| 3300001 | 3305000 | 5000 | 54347 | 2450001 | 2455000 | 5000 | 904347 |
| 1850001 | 1860000 | 10000 | 1502347 | 500001 | 510000 | 10000 | 1123472 |
| 2600001 | 2610000 | 10000 | 752347 | 1240001 | 1250000 | 10000 | 1873472 |
| 2800001 | 2810000 | 10000 | 552347 | 3500001 | 3510000 | 10000 | 147277 |
| 230001 | 280000 | 50000 | 845972 | 800001 | 850000 | 50000 | 1445972 |
| 1500001 | 1550000 | 50000 | 1829847 | 2100001 | 2150000 | 50000 | 1129847 |
| 3700001 | 3750000 | 50000 | 369777 | 3110001 | 3160000 | 50000 | 229847 |

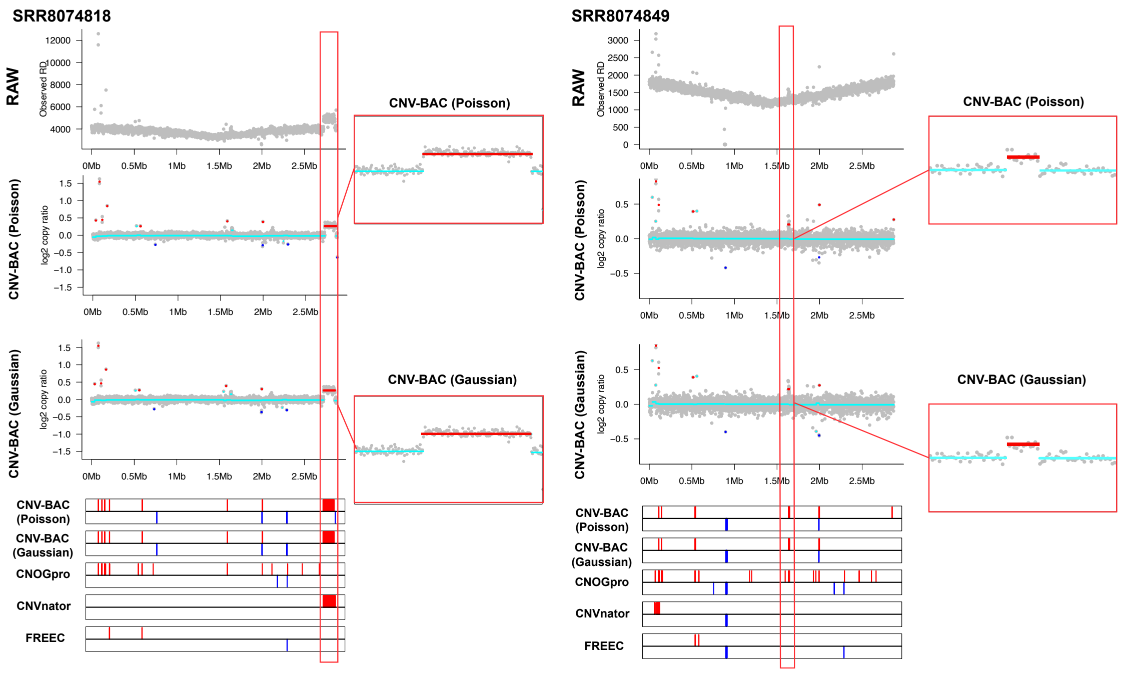


Figure S1: The read depth (RD) profile and detected CNVs of S. aureus SRR8074818 and SRR8074849. The scatter plots show the observed read counts and log2 copy ratios given by CNV-BAC, where cyan lines represent the log2 copy ratio of normal segments and red/blue lines show the log2 copy ratio for duplication/deletion segments. The bottom plots show CNVs detected by different methods, where the red and blue bars represent duplications and deletions respectively.

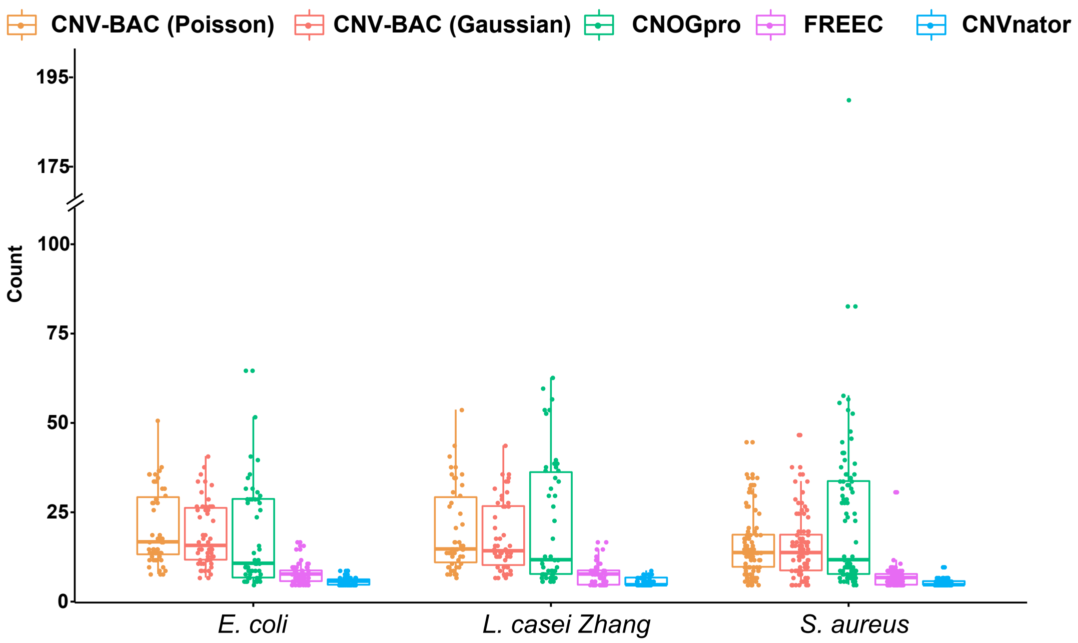


Figure S2: Number of CNVs detected by different methods in three datasets.

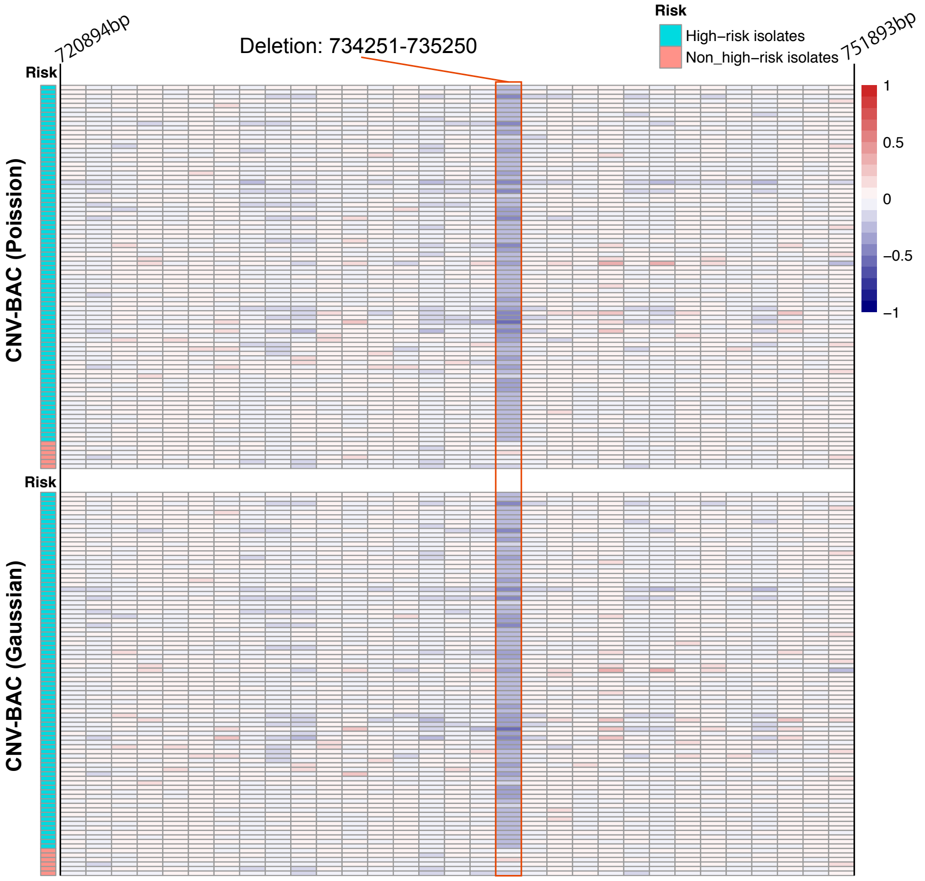


Figure S3: Heatmap of corrected log2 copy ratio for regions near deletion 734251-735250 bp in S. aureus dataset. Comparing to non-high-risk samples, most high-risk samples have low copy ratios in this region.

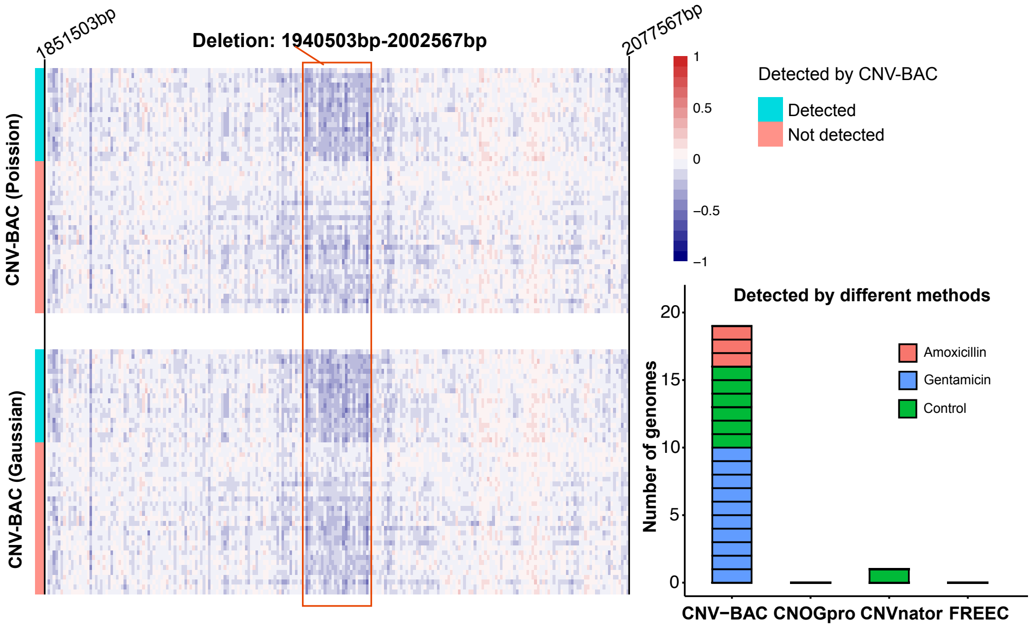


Figure S4: Deletion from 1940503bp to 2002567bp in L. casei Zhang dataset. (Left) Heatmap of corrected log2 copy ratio. (Right) Number of genomes having deletion from 1940503bp to 2002567bp detected by different methods.

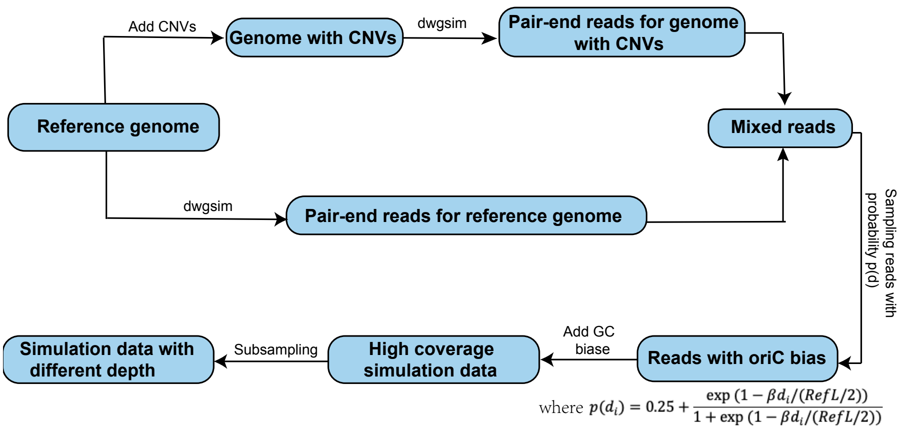


Figure S5: Pipeline for simulation sequencing data.

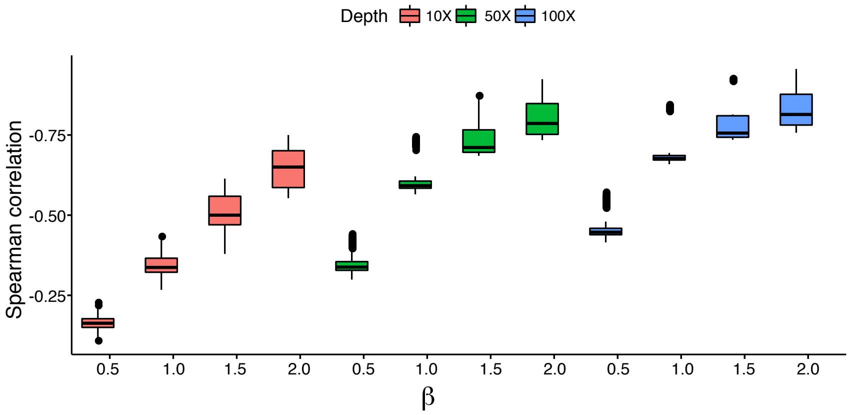


Figure S6: The Spearman's rank correlations between log2 copy ratio and distance to replication origin in simulation data.

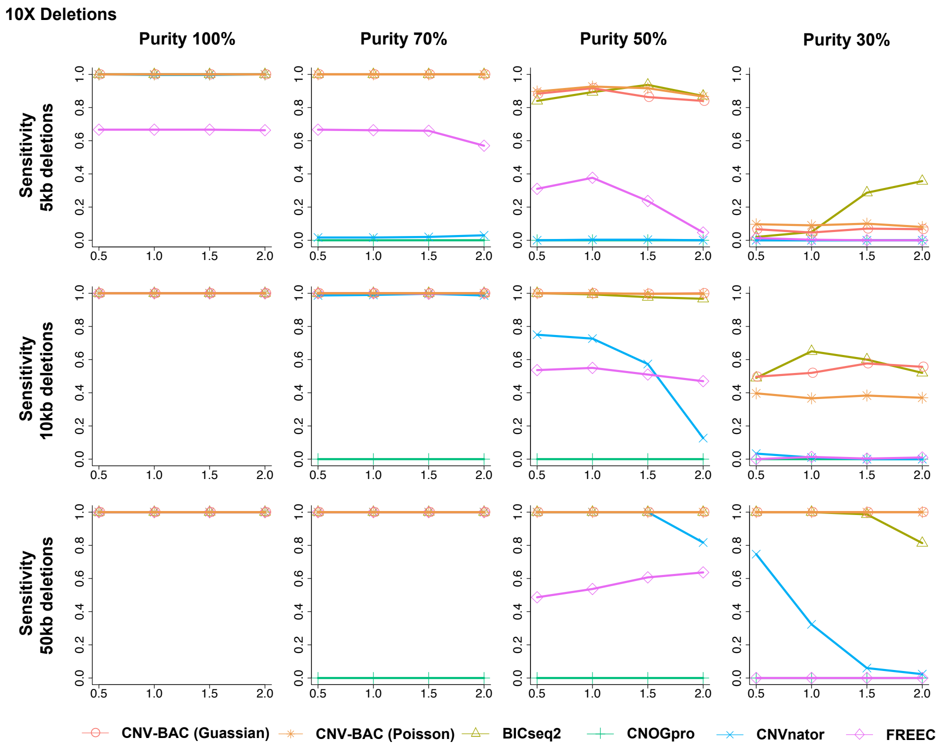


Figure S7: Average sensitivity for detecting deletions for 10X depth simulation data. The x-axis is the value of $\beta$.

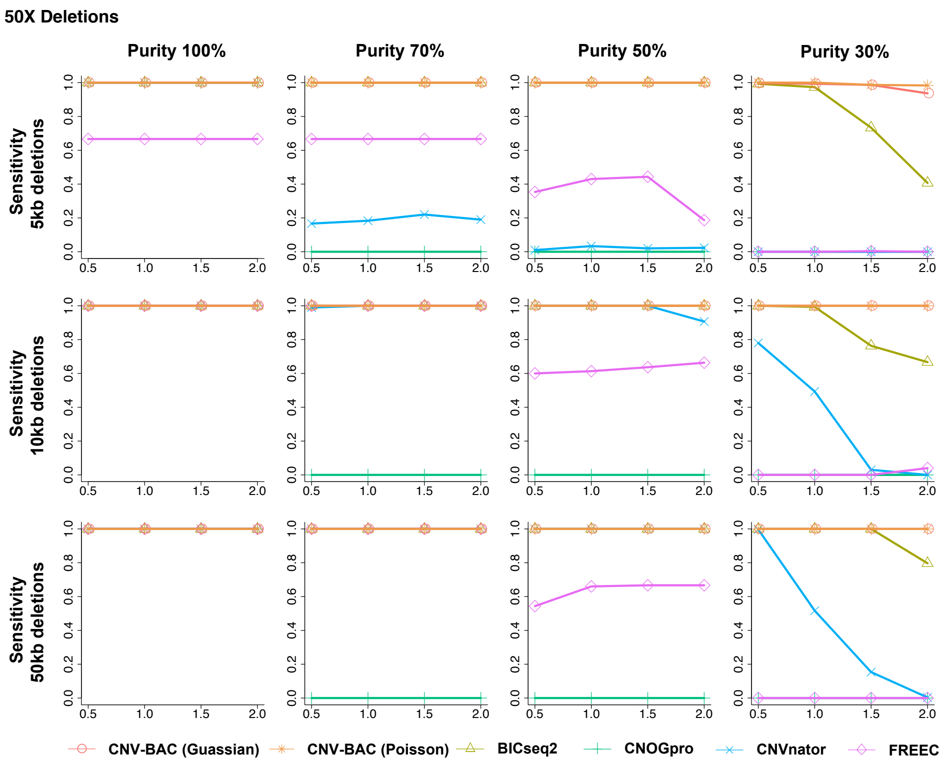


Figure S8: Average sensitivity for detecting deletions for 50X depth simulation data. The x-axis is the value of $\beta$.

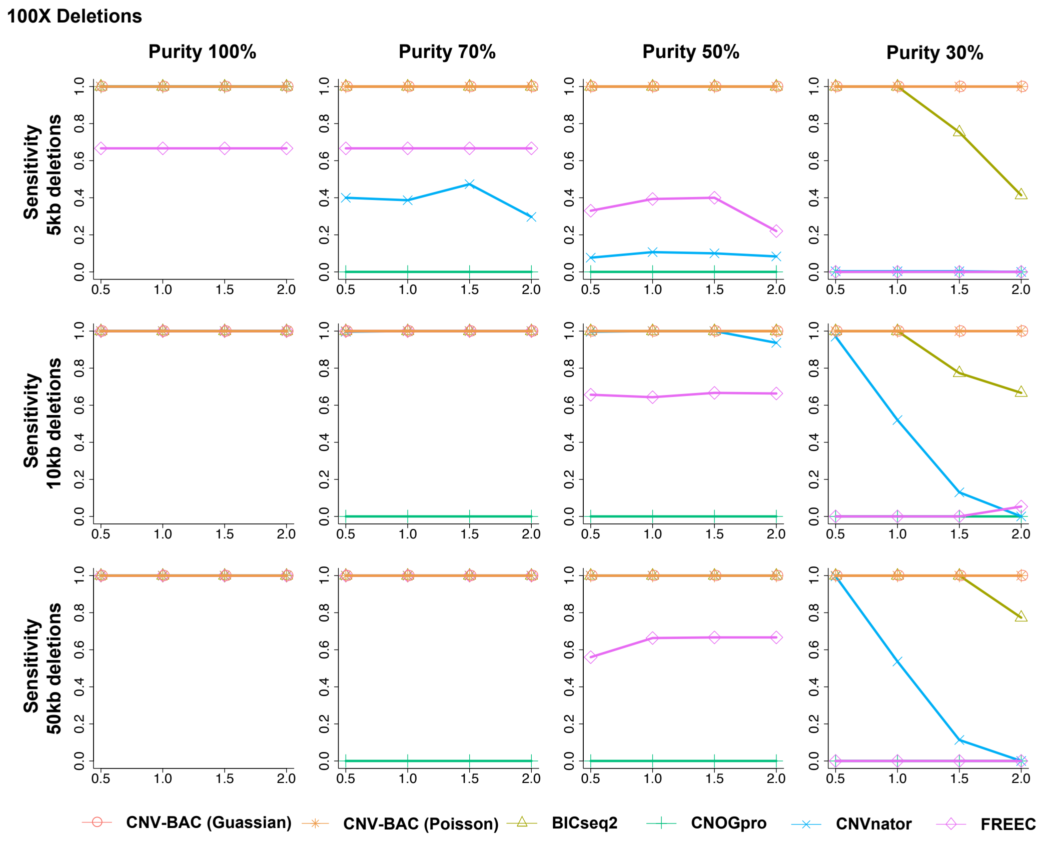


Figure S9: Average sensitivity for detecting deletions for 100X depth simulation data. The x-axis is the value of $\beta$.

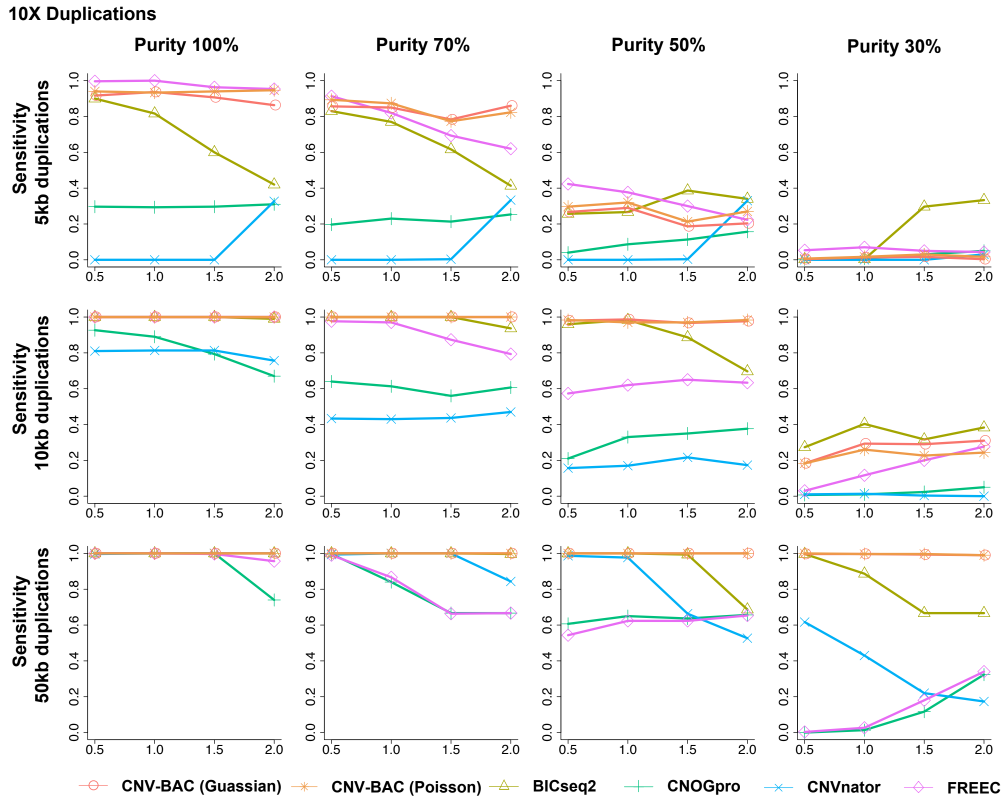


Figure S10: Average sensitivity for detecting duplications for 10X depth simulation data. The x-axis is the value of $\beta$.

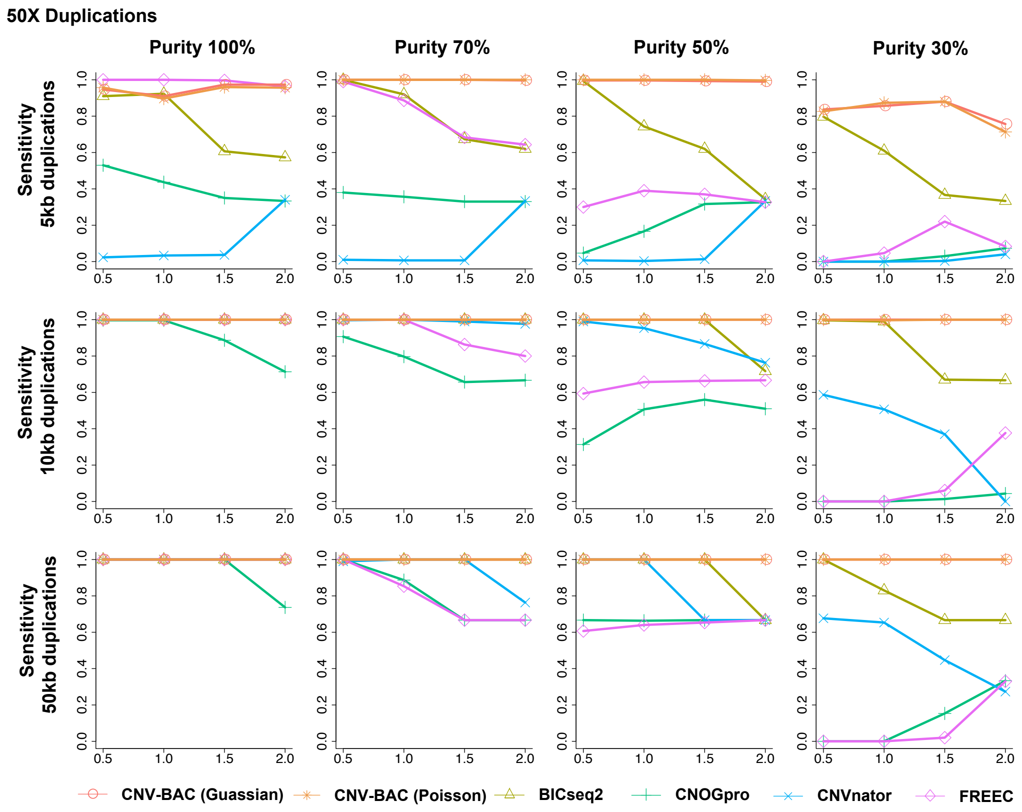


Figure S11: Average sensitivity for detecting duplications for 50X depth simulation data. The x-axis is the value of $\beta$.

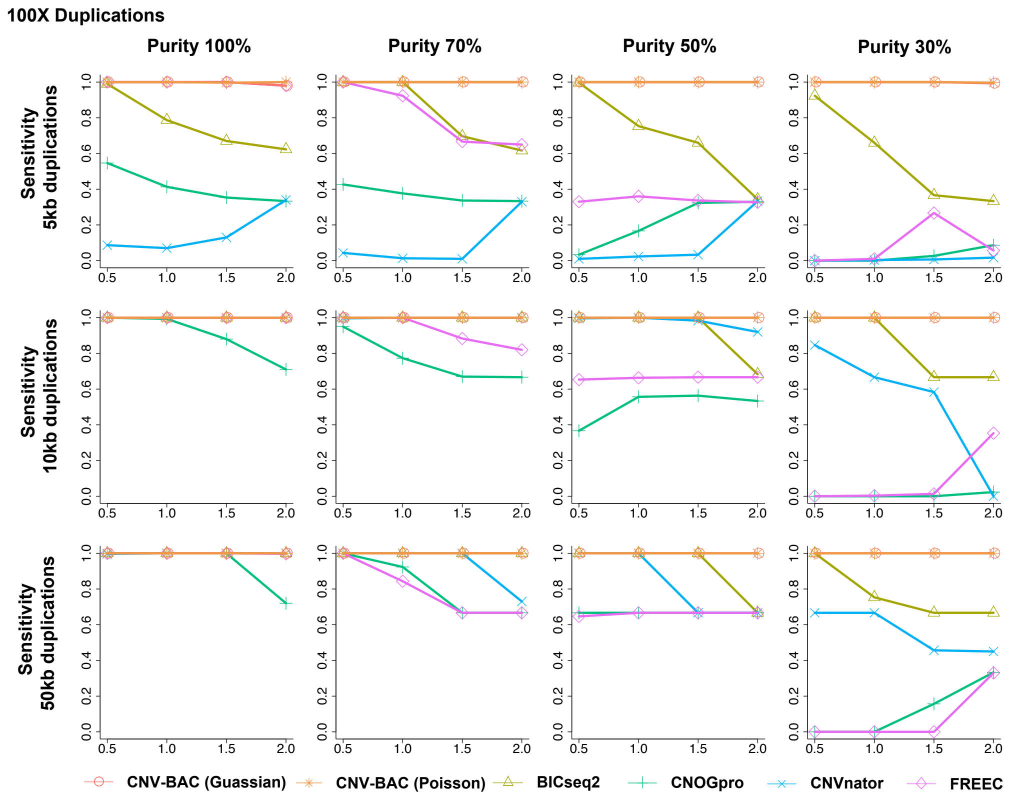


Figure S12: Average sensitivity for detecting duplications for 100X depth simulation data. The x-axis is the value of $\beta$.

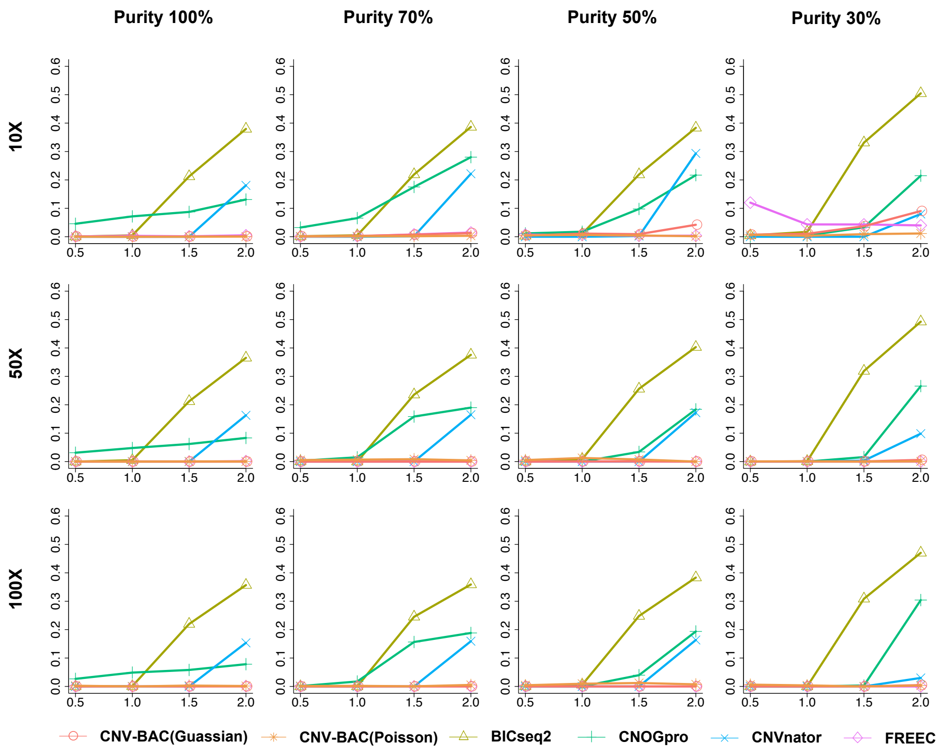


Figure S13: Average false discovery rates for simulation data. The x-axis is the value of $\beta$.

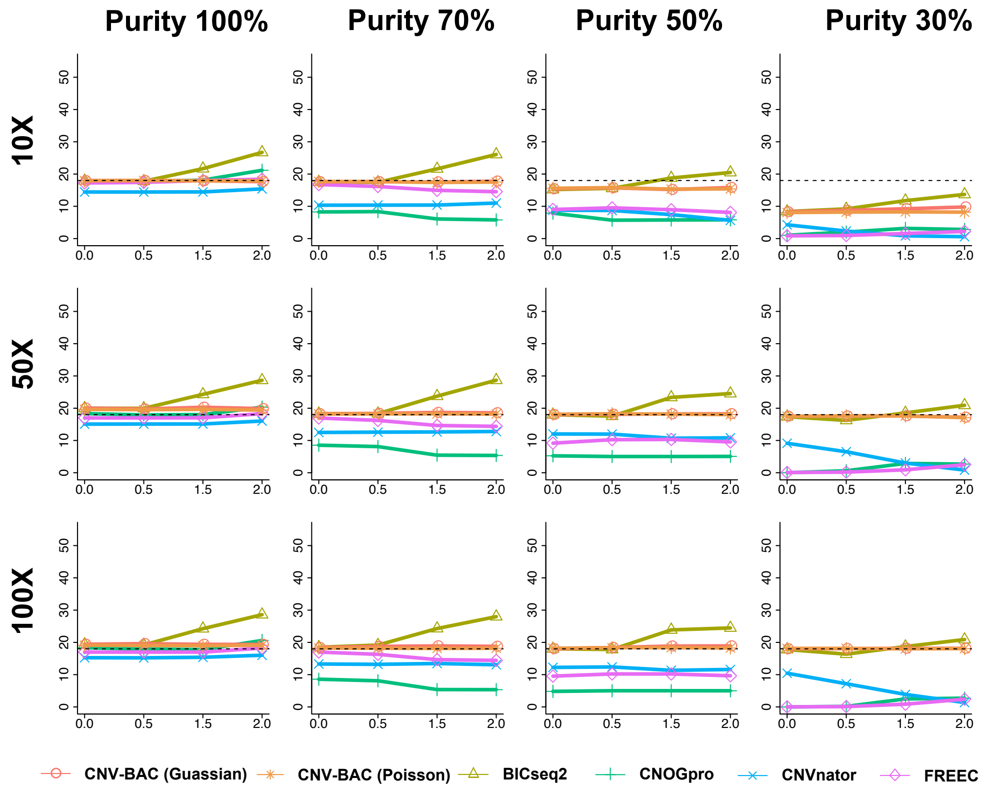


Figure S14: Average number of detected CNVs for each method in simulation. The x-axis is the value of $\beta$, and the dash line represent the number of designed CNVs.

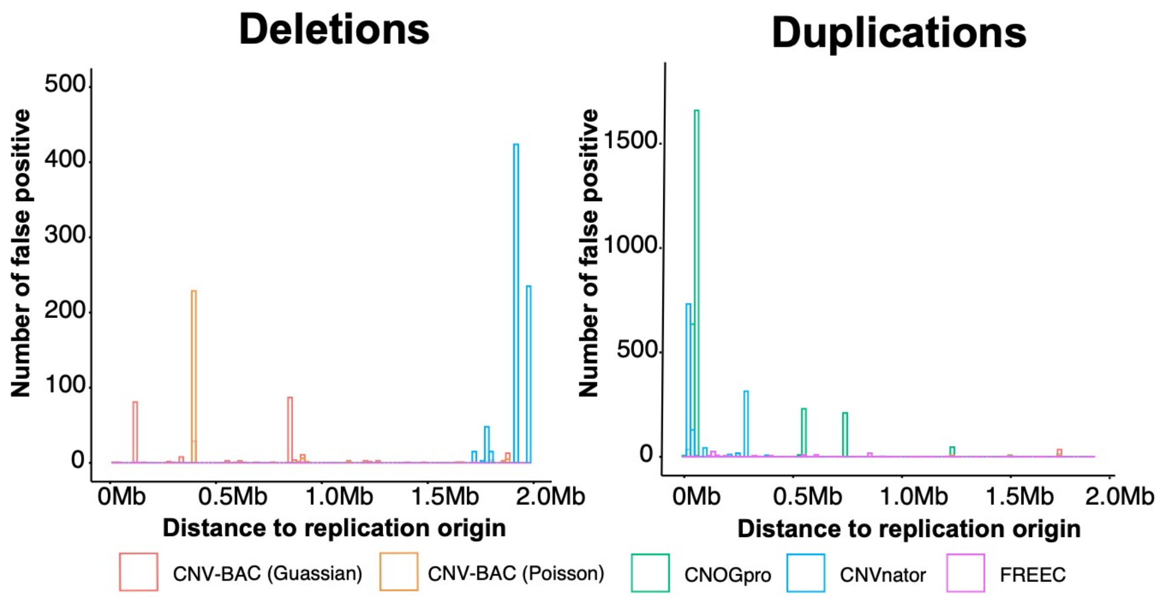


Figure S15: The distribution of distance between replication origin and the false discovered CNVs in simulation data. The x-axis is distance between replication origin and CNVs.

### Reference


Li, H. and Durbin, R. Fast and accurate short read alignment with Burrows-Wheeler transform. *Bioinformatics* 2009;25(14):1754-1760.

Luo, H. and Gao, F. DoriC 10.0: an updated database of replication origins in prokaryotic genomes including chromosomes and plasmids. *Nucleic Acids Res* 2019;47(D1):D74-D77.

Xi, R.B.*, et al.* Copy number analysis of whole-genome data using BIC-seq2 and its application to detection of cancer susceptibility variants. *Nucleic Acids Res* 2016;44(13):6274-6286.
